# Foundations of visual form selectivity in macaque areas V1 and V2

**DOI:** 10.1101/2024.03.04.583307

**Authors:** Timothy D. Oleskiw, Justin D. Lieber, Eero P. Simoncelli, J. Anthony Movshon

## Abstract

Neurons early in the primate visual cortical pathway generate responses by combining signals from other neurons: some from downstream areas, some from within the same area, and others from areas upstream. Here we develop a model that selectively combines afferents derived from a population model of V1 cells. We use this model to account for responses we recorded of both V1 and V2 neurons in awake fixating macaque monkeys to stimuli composed of a sparse collection of locally oriented features (“droplets”) designed to drive subsets of V1 neurons. The first stage computes the rectified responses of a fixed population of oriented filters at different scales that cover the visual field. The second stage computes a weighted combination of these first-stage responses, followed by a final nonlinearity, with parameters optimized to fit data from physiological recordings and constrained to encourage sparsity and locality. The fitted model accounts for the responses of both V1 and V2 neurons, capturing an average of 43% of the explainable variance for V1 and 38% for V2. The models fitted to droplet recordings predict responses to classical stimuli, such as gratings of different orientations and spatial frequencies, as well as to textures of different spectral content, which are known to be especially effective in driving V2. The models are less effective, however, at capturing the selectivity of responses to textures that include naturalistic image statistics. The pattern of afferents — defined by their weights over the 4 dimensions of spatial position, orientation, and spatial frequency — provides a common and interpretable characterization of the origin of many neuronal response properties in the early visual cortex.

## 1 Introduction

In the primate visual cortex, complex patterns are processed by a cascade of areas collectively known as the ventral stream [9, 32, 54]. At the beginning of this cascade is primary visual cortex (V1), where neurons respond selectively to spatially localized patterns of a specific orientation and scale [21]. Classically, these cells have been broadly split into two groups, known as *simple* and *complex*. Simple cells respond selectively to the local phase of an oriented pattern, and are well-described by a linear-nonlinear (LN) model: a single spatial filter followed by a rectification step [17, 22, 34]. Complex cells respond independently of local phase, and are well-described by the squared sum of two spatial filters, a construction known as the “energy model” [1, 33]. In practice, many cells in V1 lie on a continuum between these two extremes [30, 37], and the full set can be described by a unified model that spatially pools local populations of rectified linear filters [56]. These models, when combined with divisive normalization and contextual modulation by the receptive field surround, are state-of-the-art in predicting the responses of V1 neurons to natural images [5, 42].

Extrastriate visual area V2, directly downstream of V1, has been implicated in the processing of visual patterns that are more complex than simple oriented lines. Like neurons in V1, many in V2 respond selectively to spatially localized patterns of specific orientations and spatial frequencies [24]. However, V2 responses are often wider in orientation and spatial frequency bandwidth than those in V1, as would be expected if V2 neurons were pooling the responses of multiple V1 inputs [24]. Moreover, lesions in V2 produce deficits in higher-order feature detection while preserving orientation perception [31], suggesting a specific, causal role for V2 in the perception of complex form. When probed with more complex stimuli, some V2 neurons respond selectively to curved contours [2, 18], object boundaries[61], and multi-point correlations in binary texture patterns [60]. Finally, many neurons in V2 respond selectively to the higher-order statistics of “naturalistic” textures, more so than neurons in V1 [12].

How can one construct a model to characterize the diverse form-selectivity of V2 responses? It is generally assumed that visual pattern recognition emerges through cascaded transformations of canonical computations [13, 15], with an initial stage corresponding to area V1. These multi-stage models can explain perceptual phenomena, such as second-order texture boundary detection [4, 10, 57], and are, in simplified form, the substrate for deep neural network models developed in the field of machine learning for object recognition [23].

Cascade models have been used to fit the responses of neurons in V2 [26, 40, 51, 58]. However, several factors make these fits difficult to interpret. First, many of these studies use images of natural scenes [26, 40, 58], which are highly correlated in the parameter space of these models. While stimulus bias can, in principle, be addressed by incorporating a regularizer or prior distribution over the parameter space, in practice, this requires many more presentation trials than can be collected in an awake experimental preparation. Second, these approaches have used rapid sequential stimulus presentations to estimate a neuron’s receptive field in space and time [7, 35]. Given the dynamic selectivity of V1 neurons [29, 39], particularly over the extra-classical receptive field [6, 19, 43, 50, 59], it is unclear whether V2 response patterns measured using rapidly changing stimuli will extrapolate to more natural viewing conditions.

In this study, we develop an interpretable account of neuronal receptive fields in macaque V1 and V2. We designed a set of stimuli – sparse flashed localized oriented elements, termed *droplets* – that reliably activate V1 and V2 neurons while possessing limited structural regularities in image space. We presented these stimuli at an ethologically relevant cadence, roughly matched to typical primate inter-saccade intervals [3]. Droplets allow for a robust and (relatively) unbiased model fitting of early ventral visual neurons by foregoing the complex statistical dependencies of natural images. We fit neurons using a sparse combination of a fixed set of rectified filters that are selective for orientation and spatial frequency. Neurons in both V1 and V2 were well driven by droplet stimuli and well fit by the model. Neurons in V1 were more tightly tuned in orientation and spatial frequency. In V2, but not V1, we found a subpopulation of neurons that combine different orientations and are selective for more complex image features.

## 2 Methods

### 2.1 Stimulus design

A sequence of visual stimuli was presented for brief (200ms) intervals, separated by equal-duration blank intervals. Each stimulus comprised a randomly selected set of windowed sinusoidal gratings, hereafter “droplets”. The window centers were arranged on three hexagonal lattices, each covering the same spatial extent, with three, five, and seven vertices in diameter (containing 7, 19, and 37 windows, respectively - Fig. 1a). For each lattice, the associated windows were circular, with raised cosine transitions at the boundaries of radii chosen to allow smooth coverage of the full extent of the stimulus. Lattice size was chosen so that the central window of the coarsest lattice matched the classical receptive field size of the recorded unit (Fig. 1c).

**Figure 1:**
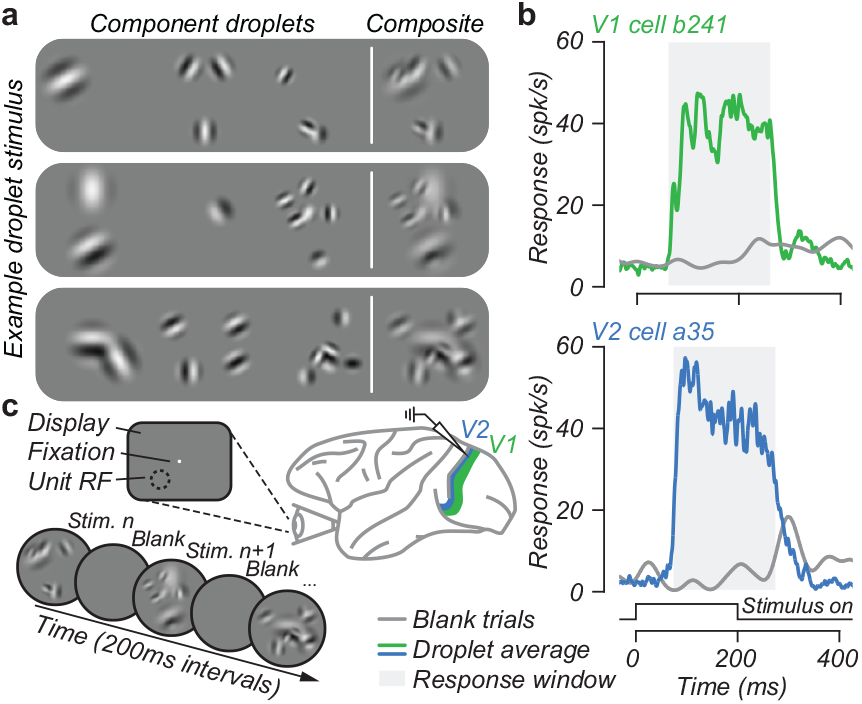
Stimulus design and recording paradigm. (a)Three example stimuli, each constructed as a sum of windowed grating elements (“droplets”) randomly selected from a set spanning a range of locations, spatial frequencies, orientations, and phases (see text). (b) Average peristimulus time histogram of exemplar V1 and V2 neurons to droplet stimuli, compared to its baseline response (blank trials, gray line). (c) Experimental design. A sequence of randomly selected droplet stimuli are presented for 200ms intervals, separated by 200ms of blank screen.

For each window, a single sinusoidal grating was randomly selected from a set of 48, covering six orientations (including horizontal and vertical), two different spatial periods (differing by a factor of two and scaled according to window size), and four spatial phases (odd- or even-symmetric). The base frequency (period) of the gratings was hand-adjusted to evoke strong responses from the unit. Finally, a subset of windows within each lattice was randomly selected for inclusion in the stimulus, with a probability of 20% (the others were turned off). The set of selected gratings was multiplied by their associated windows and summed, and the range of this image was then clipped to lie within the display’s dynamic range.

In addition, for comparison with model predictions, we measured responses to static flashed grating and texture targets. Grating targets were presented within a window matched to the hand-mapped receptive field at 8 uniformly-spaced orientations and 5 logarithmically-spaced spatial frequencies (0.25 − 4c/deg). Each grating’s phase was chosen randomly per presentation, either odd or even symmetric. The texture stimuli were those described in [63] and comprised 16 instances of 5 naturalistic texture families and their spectrally-matched (noise) controls presented as a circular patch 4deg in diameter.

### 2.2 Neuronal recordings

We collected neurophysiological data from areas V1 and V2 of two adult male rhesus macaques trained to fixate a small target for fluid reward. Before recording, we implanted a custom headpost for head stabilization using a standard design and methods described previously (Grey Matter Research, Bozeman, MT). In a subsequent surgical procedure, we implanted a recording chamber over the Lunate Sulcus of the right hemisphere. Chamber placement and design were guided with structural magnetic resonance imaging (MRI) and visualization software (Brainsight, Rogue Research) to produce a chamber with stabilizing legs that matched skull curvature. All procedures complied with the National Institute of Health Guide for the Care and Use of Laboratory Animals, with the approval of the New York University Animal Welfare Committee.

We acclimated each monkey to the recording chair and experimental surroundings. After this initial period, the animals were head-restrained with their eyes 57cm from the display and rewarded with diluted juice or water for fixating a target. Meanwhile, we used an infrared eye tracker (EyeLink 1000; SR Research) to monitor eye position via reflections of infrared light on the cornea and pupil at a sampling rate of 1 kHz.

Stimulus presentation and time sequence are illustrated in Fig. 1. The monkey initiated a trial by fixating on a small white spot (diameter 0.1 − 0.2deg), after which he was required to maintain fixation for a 200 − 500ms interval. Stimuli were presented for 200ms, followed by a 200ms inter-stimulus blank screen interval (Fig. 1c). The monkey was rewarded for maintaining a gaze signal within 1 − 1.75deg of the fixation point for 4 to 8 presentations. After accounting for estimation noise, the center of gaze was typically within 0.5deg of fixation during valid presentation trials. No stimuli were presented during the 300 − 600ms in which the reward was delivered. If the monkey broke fixation prematurely, the trial was aborted, no reward was given, and a timeout of 1 *−* 2s was imposed.

Recordings were conducted by advancing a 6 − 10MΩ impedance tungsten-epoxy microelectrode (FHC) through a 23 Gauge stainless steel guide tube that was stabilized with a customized 3D-printed grid insert affixed to the recording chamber. Both the grid and guide tube were held in contact with the dura during electrode penetration. We distinguished V2 from V1 based on depth from the cortical surface and changes in the receptive field location of recorded units. To obtain an unbiased sample of single units, we made extracellular recordings from every single unit with a spike waveform that rose sufficiently above background noise to be isolated. Data are reported from every unit for which we completed characterization. The receptive fields of most units were centered 1 *−* 5 deg from the center of gaze, estimated via manual RF characterization.

After isolating a single-unit spike waveform, we estimated receptive field location and size with hand-positioned targets. We estimated orientation and spatial frequency tuning by presenting sinusoidal gratings randomly interleaved at 200ms on/off intervals (see stimulus design). The primary parameters of the droplet stimuli – their size and spatial frequency – were then set to be consistent with the estimated receptive field size and spatial frequency preference.

Neural response to each stimulus was determined from the mean evoked activity, estimated from each unit’s peristimulus time histogram (PSTH) computed for stimulus-present and stimulus-absent (baseline) trials (Fig. 1b). Mean responses were computed by averaging the number of spikes within a time window of duration matching that of the stimulus presentation (200 ms), beginning at the unit’s onset response latency.

### 2.3 A two-stage canonical network model

To capture the visual response properties of each recorded neuron, we use a two-stage network (Fig. 2), in which units of each stage consist of a spatially localized linear weighted sum of inputs, followed by a rectifying nonlinearity (we refer to each stage as “LN”). The first “afferent” stage implements a stylized population of V1-like units, whereas the second stage implements a single unit whose parameters are optimized for each individually recorded neuron. Similar “convolutional subunit” models have been used to describe responses in retina [11, 25, 28, 44, 45, 55] and cortical areas V1[14, 38, 42, 46, 56], MT[41, 48], and V2[26, 40, 51, 58, 62].

**Figure 2:**
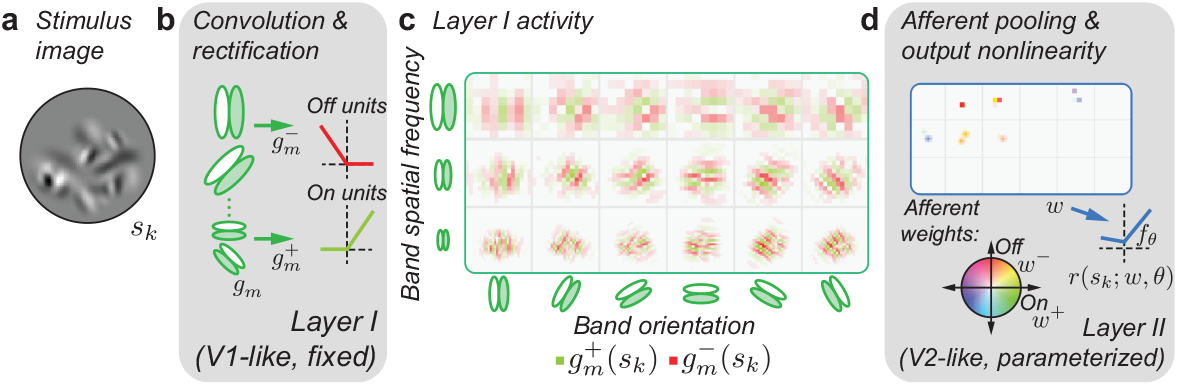
Visualization of the two-stage model stimulus response. (a) An example droplet stimulus. (b) Stage one is constructed by convolution of the stimulus with a fixed bank of band-pass oriented linear filters, followed by (positive and negative) half-wave rectification. (c) Stage one responses tile visual space, mimicking a population of V1 simple cells. (d) Stage two consists of a linear combination over stage one activity, followed by a piecewise-linear two-sided rectifier.

The linear portion of the first stage is computed by spatial convolution with a predetermined bank of 36 oriented band-pass filters, known collectively as a “steerable pyramid” [47]. Individual filters have bandwidths of one octave and orientation responses covering 60 degrees, approximating the selectivity of a typical V1 neuron. Filters have peak spatial frequency responses at three frequencies (differing by factors of two) and six orientations (at intervals of *π/*6 radians). They also come in even and odd-symmetric form. Spatial frequency and orientation selectivity are precisely chosen so that the Fourier spectra of the full set of filters provide uniform coverage of the full range of orientations and three octaves of spatial frequency. The responses of the full set of linear filters constitute a complete and invertible representation of the image. More importantly, for our purposes, the full set of filters can be considered as a stylized population of V1-like units (Fig. 1d). We denote the channels resulting from the convolution of the *k*^th^ stimulus image, *s*_*k*_, with the *m*^th^ filter as *g*_*m*_(*s*_*k*_). To mimic V1 simple cell responses, these linear responses are rectified and partitioned into the positive and negative portions:

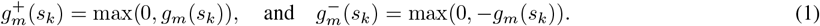

The second stage of the model computes a weighted sum of the first-stage responses, followed by another nonlinearity, generating the firing rate response

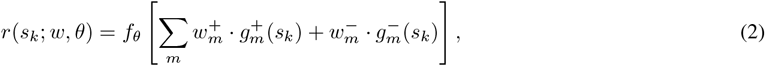

where *⋅* indicates a dot product between the weights and responses, and the parameter vector *w* contains the full set of real-valued weights (corresponding to positive and negative parts of responses within all 18 first-stage channels, 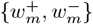). The function *f*_*θ*_() implements a final nonlinearity,

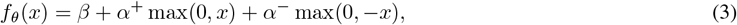

with parameters *θ* = {*α*^+^, *α*^*−*^, *β*} encoding the positive gain, negative gain, and offset, respectively.

### 2.4 Model optimization and regularization

To train the model we seek the second-stage weights *w* and affine response parameters *θ* that minimize the normalized squared error between measured responses *r*_*k*_ (spike count, in response to the *k*^*th*^ stimulus *s*_*k*_) and corresponding model predictions:

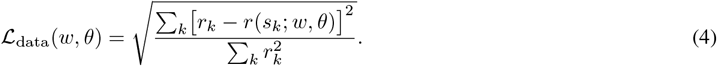

Since the number of model parameters is typically much larger than the number of unique stimuli presented, the optimization problem is underdetermined and susceptible to over-fitting. To address this, we incorporated two regularization terms that bias the weights *w* toward *sparsity* and *locality*. Sparsity is induced on the pairs of signed weights. Specifically, we define weight amplitudes as

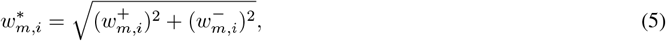

where *i* is an index for the spatial location. Sparsity is induced on these amplitudes by penalizing the effective dimensionality of *w*^*∗*^ (*L*_1_-norm to *L*_2_-norm ratio), which provides a continuous approximation of the square root of the number of non-zero elements:

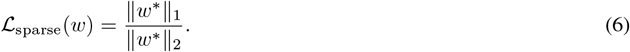

The normalization ensures that the regularization penalty does not drive the vector of weights toward zero, compensating with increases in second-stage parameters {*α*^+^, *α*^*−*^}. The sparsity penalty is imposed independently on weights at each position and channel without regard to their proximity. To encourage smooth and contiguous afferent maps, we also included a regularization term to penalize spatial and spectral dispersion of coefficient magnitudes. Specifically, we computed the spatial variance of weight maps in the horizontal (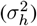) and vertical (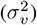) directions, and the spectral variance across channels in orientation (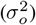) and scale (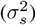). For example, the horizontal variance is computed as:

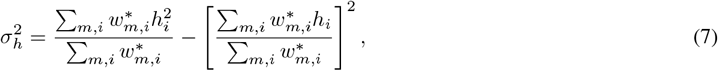

where *h*_*i*_ indicates the horizontal position associated spatial location *i*. Note that, as with the sparsity term, these localization terms are invariant to the scaling of weight vector *w*, ensuring that an overall reduction of weight amplitudes is not encouraged. Spectral variances are in octaves, and spatial coordinates are scaled such that an octave represents twice the estimated receptive field diameter to equilibrate the units of these variances. The locality regularizer is then expressed as

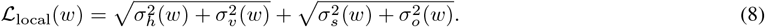

For each neuron, model fitting proceeds by minimizing the weighted sum of the data fidelity objective and the two regularization terms:

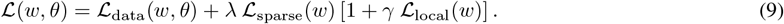

The locality bias parameter *γ* is empirically chosen to penalize afferent dispersion at approximately 1% the magnitude of the coefficient sparsity term. The regularization parameter *λ* is chosen for each cell using a cross-validation procedure. Specifically, stimulus-response pairs are randomly assigned to one of ten equal-sized partitions. Each partition is held aside as a test set, the model is optimized over the stimulus-response pairs from the other nine partitions, and a test error is then computed for the data in the test set. The value of *λ* is optimized to minimize error on the held-out test sets. Training (fitting) and testing errors are computed from the sum of squared error between observed responses and model predictions, normalized by the variance of observed responses.

## 3 Results

### 3.1 The model explains both V1 and V2 responses

We quantify how well our two-stage model explains V1 and V2 activity by plotting the variance explained across training and testing partitions. Fig. 3a,c shows training performance as a function of testing performance - their similarity indicates the model is not over-fit. A fraction of the cells in our population were poorly fit by the model (V1:*n* = 16 of 69, V2: *n* = 28 of 120), exhibiting very low training (*<* 0.1) performance. Upon inspection, these units either had insufficient recording time or insufficient evoked activity to support effective model fitting. Some of these units were also most active during stimulus offset, which was not included in our response activity window and thus was not accounted for in our analysis.

**Figure 3:**
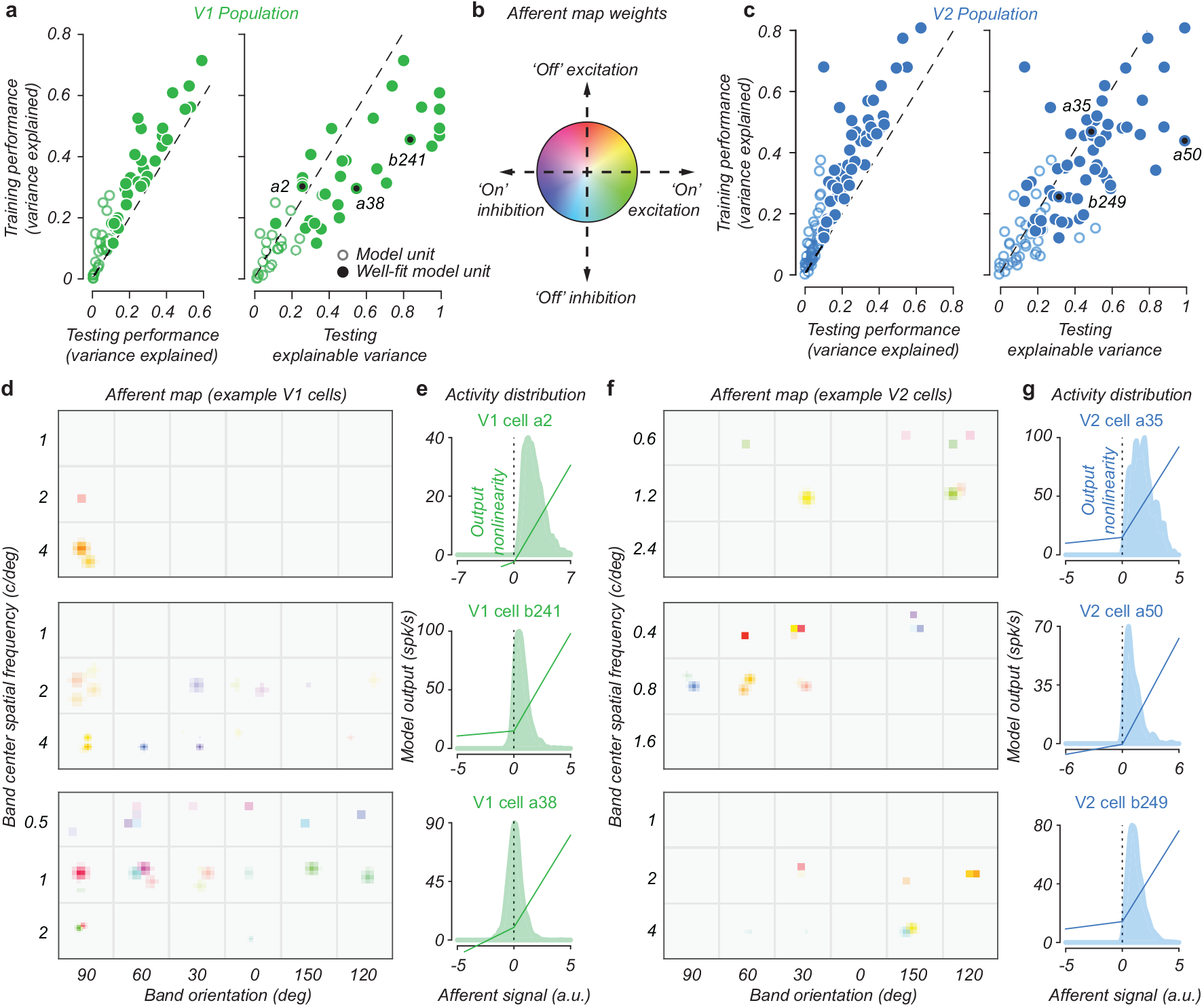
Model performance and example unit fit. Training and testing performance of models fit to (a) V1 and (c) V2 neurons in our population. (b) Each model optimized the connection weights projecting from a V1-like afferent representation. The hue and saturation of afferent connections convey the proportion of on (*w*^+^) versus off (*w*^*−*^) weights and their amplitude (*w*^*∗*^), respectively. Model fits of three V1 and three V2 cells are visualized by their afferent maps (d,f) and output nonlinearities (e,g), respectively.

Of the remaining units, not all generalize well, in some cases due to high neuronal variability. We judge that these fitted models are unlikely to reflect true neuronal tuning characteristics. Therefore, for the balance of the paper, we analyze the properties of those V1 and V2 units with testing V.E. *>*= 0.1. These “well-fit” model units (Fig. 3a,c, solid symbols) comprise 32 of the 69 V1 units (46%) and 56 of the 120 V2 units (47%).

The variance explained by these “well-fit” units is limited by intrinsic neural variability. To measure the amount of *explainable variance, i*.*e*. the predictive power of a model up to independent neuronal variability, we first estimate the noise ceiling of each recording. For each dataset, we identified stimuli with three or more repeated trials, computing the mean and variance of the response to each stimulus. To model trial-to-trial variability, we assume neuronal response variance is proportional to its mean and find the proportionality factor that minimizes the squared error between mean and variance pairs across all stimuli for a neuron. Then, we simulate a *synthetic* recording session, sampling firing rates under the fitted noise model for every recorded trial. We simulate repeated stimuli to sample neuronal activity from a set of trials matched to those recorded. This process is repeated and compared against observed responses to estimate an upper bound for the portion of response variance that can be explained by *any* model. By normalizing the testing variance explained by this noise ceiling, we plot in Fig. 3a,c (right-hand panels) this explainable variance. Many neurons generalize well to capture a considerable fraction of the stimulus-response variance expected to be explained by a model. Of units that were sufficiently driven by our stimulus for the model to converge (training V.E. *>* 0.1), the average explainable variance captured was substantial (V1: 43%, *n* = 53; V2: 38% *n* = 92).

To visualize optimized model parameters, we show in Fig. 3d a representation of the connection weights *w* and output nonlinearity *f*_*θ*_ of 3 V1 and 3 V2 example units. Connection weights are grouped into spectral bands of orientation (columns) and spatial frequency (rows) as in Fig. 2. Within each band, weights are arrayed in their natural horizontal and vertical spatial position, *i*.*e*. each band represents the same region of visual space. We superimpose the two independent rectified channels for each spectral band, using saturation to denote absolute connection magnitude and hue to encode the relative contribution of each channel. We denote this representation of projection weights, pooling contrast across spectral and spatial domains, the *afferent map*.

### 3.2 Linearity and separability of afferent maps

The afferent map of connection weights, representing a linear combination of basis elements tuned in orientation and spatial frequency that tile visual space, is segregated into positively and negatively rectified terms. However, given that afferents are linearly combined, we can rotate the basis to organize afferent activity along more interpretable axes, corresponding to a linear response (difference of the positive and negative parts) and the absolute value (sum of the positive and negative parts, which we refer to as *energy*):

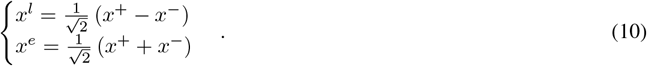

By similarly transforming the optimized connection weights *w* to yield *w*^*l*^ and *w*^*e*^, we can examine the contribution of linear and nonlinear response features in the fitted model.

Specifically, we determine the fraction of response variance explained by the afferent map attributed to the linear and energy components in isolation. To do this, we project stimulus activity *g* (*S*_*k*_) for each *k* ∈ *K* against *w*^*l*^ and *w*^*e*^, computing the squared correlation coefficient (Pearson’s *r*^2^) between recorded neuronal activity. The linear and energy component map *r*^2^ is normalized by the total afferent map *r*^2^, up to the output nonlinearity, independently for each model. This fraction of variance explained by each model’s linear and energy components is shown in Fig. 4a. Note, the total fraction variance explained is *approximately* equal to 1 due to the output nonlinearity and coefficients shared between components. We define a linearity index as the difference in variance explained by the linear and energy component maps.

**Figure 4:**
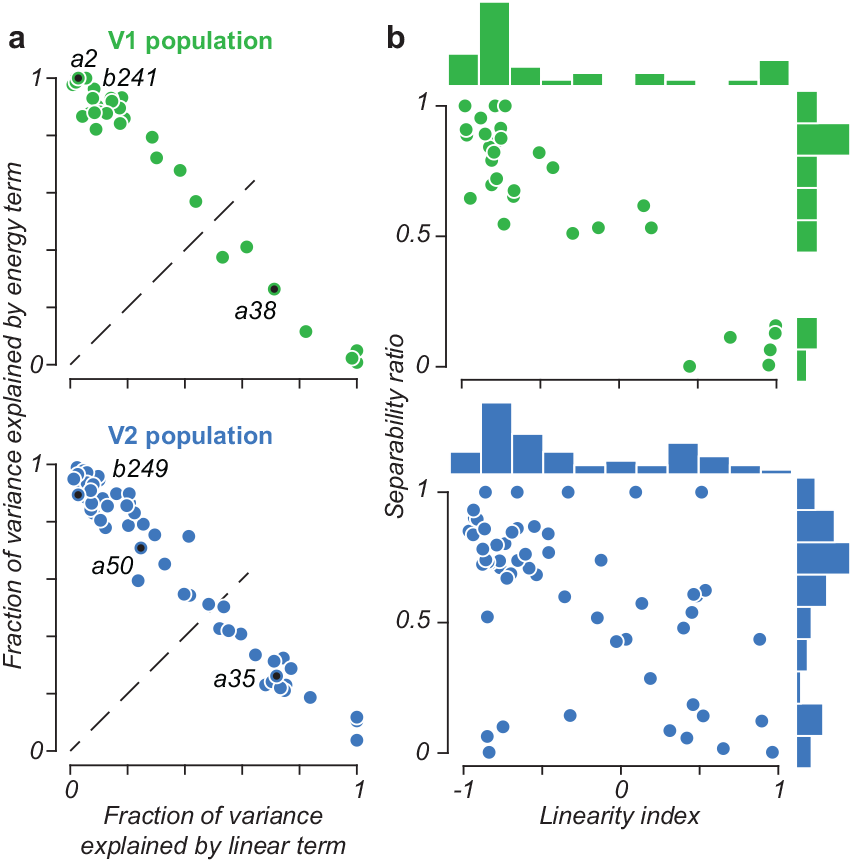
Model unit linearity and separability. (a) Both V1 and V2 populations span a continuum in terms of their construction from linear vs. non-linear (energy) afferents. (b) Linearity of afferent maps is anti-correlated with their separability along the cardinal dimensions of orientation, scale, and spatial location, especially in V1.

Next, we assess to what extent the connection weight maps are separable along the cardinal dimensions of afferent tuning. To achieve this, we approximate each afferent map with the product of three functions computed from the marginal of connectivity in 2D space, orientation, and scale. We define a separability ratio of each model unit as the fraction of variance explained by the separable afferent map relative to that of the original map. For V1 and V2 model units, we find in Fig. 4b a significant relationship between afferent separability and linearity, with linear units tending to be inseparable along afferent tuning dimensions (Pearson’s *r*; V1: *r* = .91, *p <* .0001 & V2: *r* = .52, *p <* .0001). This relationship appears most evident in V1, with no well-fit V1 model units being nonlinear and inseparable or separable and linear. It is also notable that the V2 population seems to have an over-representation of units that are both linear and non-separable (*i*.*e*. are in the bottom right quadrant). Indeed, for V1 and V2 units with separability ≤ .5, the distributions of linearity indices are significantly different (Wilcoxon rank-sum test, *p* = .007).

### 3.3 Interpreting selectivity: the afferent field

As our analysis of the afferent map shows V2 model units to be well-characterized by their constituent linear or energy components, we seek a method to interpret the spatial and spectral organization of each component beyond the afferent map representation illustrated in Fig. 5a. Since the steerable pyramid basis is a linear transformation of pixels, the linear map *w*^*l*^ of pyramid coefficients can be inverted to visualize its contribution to model response in image coordinates. We denote the visualization of this component as the model’s “linear receptive field”, analogous to that achieved via reverse correlation to characterize V1 simple cells [8, 22].

**Figure 5:**
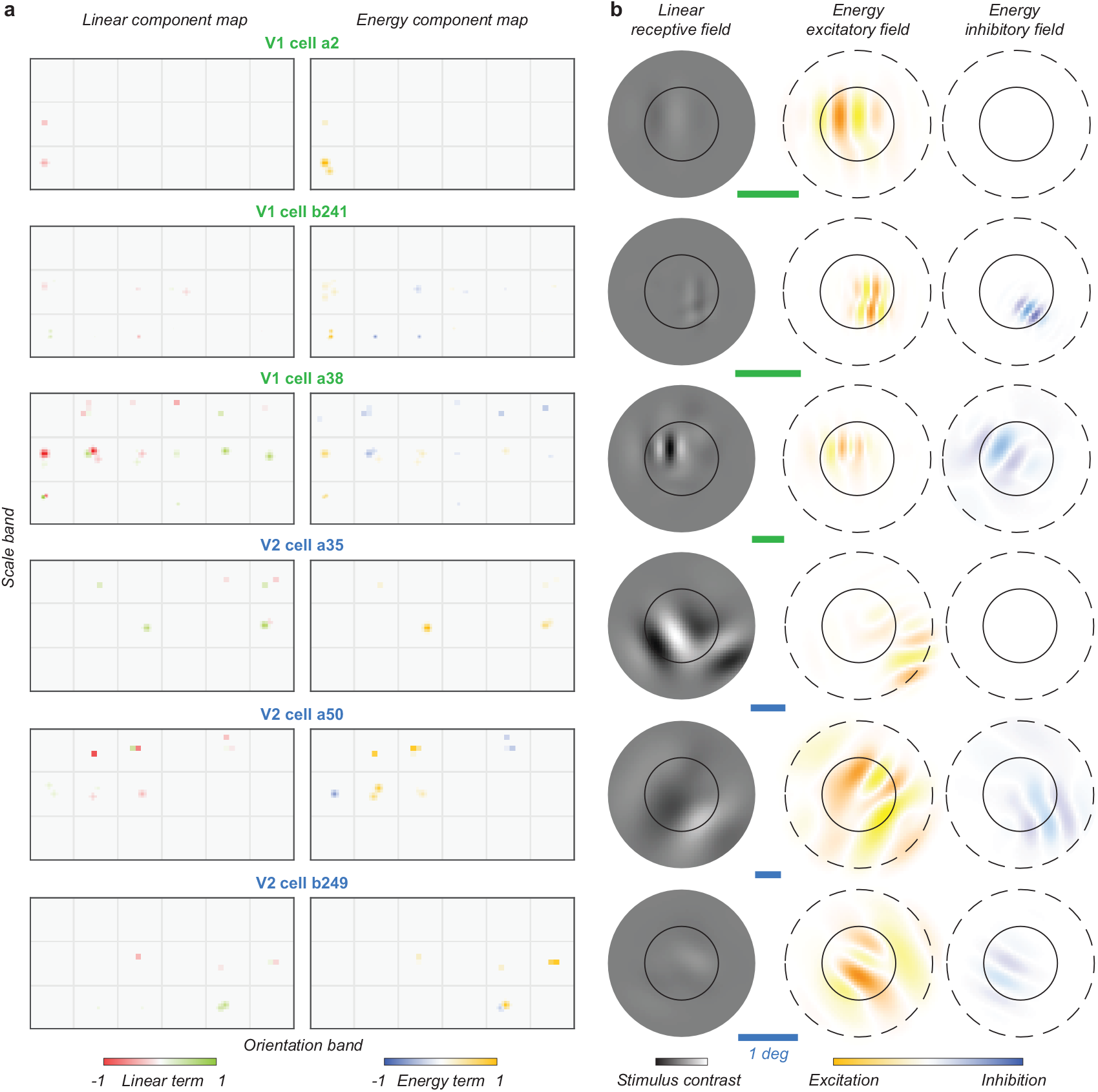
(a) Linear and energy components of example V1 and V2 afferent maps (see Fig. 3d). (b) Afferent field visualization of each unit’s selectivity. Circles overlaying the linear and energy fields denote the cell’s receptive field diameter as estimated from hand-mapping (solid) and droplet stimulus extent (dashed). Scale bars denote 1 degree of visual angle.

Interpretation of the energy component weights, however, cannot be achieved with the same visualization technique since multiple pixel representations of spectral energy exist up to the sign of all non-zero *w*^*e*^ terms. Instead, the spectral and spatial arrangement of *w*^*e*^ is depicted by producing a separate receptive field image for excitatory (*w*^*e*^ *>* 0) and inhibitory (*w*^*e*^ *<* 0) terms. We choose a characteristic image that minimizes destructive interference due to the phase (sign) of pyramid coefficients. The sinusoidal image components are shown with warm and cool hues to convey phase-invariant spectral power of the excitatory and inhibitory energy fields, respectively. Finally, the contrast of linear and energy receptive fields are scaled to convey their relative contribution to the model’s response as determined from the variance explained by *w*^*l*^ and *w*^*e*^, respectively, preserving the relative difference in magnitude between excitatory and inhibitory fields. Together, this afferent field, shown in Fig. 5b, yields a spatial representation of a model unit’s linear and energy components that depict its selectivity.

### 3.4 Spectral bandwidth of model afferents

To localize afferent connectivity, we computed the mean and variance of model weights in both orientation and spatial frequency. In Fig. 6 we plot spectral bandwidth as a function of preferred spatial frequency for well-fit units from our V1 and V2 populations. Fig. 6a,d shows our population to uniformly cover preferred orientations, with a distribution of spatial frequency coverage expected at the eccentricity of our recordings. As expected, we found a relationship between the bandwidth of spatial frequency connections and their mean (Fig. 6b,e). Conversely, we found no relationship between the mean of spatial frequency afferents and the bandwidth of orientation tuning (Fig. 6c,f).

**Figure 6:**
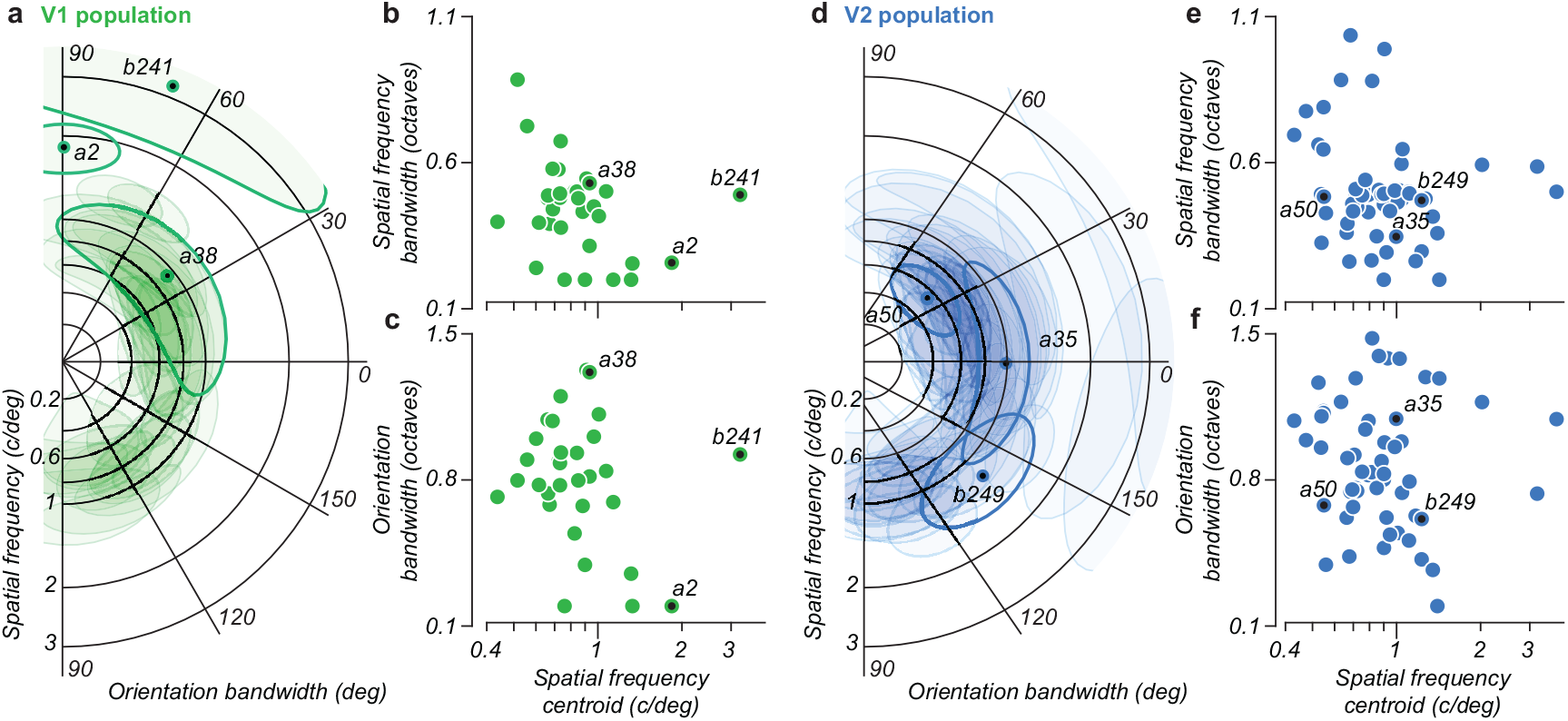
Spectral bandwidth of afferent connection weights from well-fit units from our population of (a) V1 and (d) V2 neurons. Each unit is plotted as the log-polar contour of an ellipse (shaded), with position and elongation depicting the centroid and standard deviation of afferent weights, respectively. Exemplar units are outlined for clarity. The bandwidth of (b,e) spatial frequency (c,f) and orientation is plotted versus the afferent map’s spatial frequency centroid.

An inspection between our V1 and V2 populations reveals no apparent qualitative difference between the spectral bandwidth of well-fit model afferents, with a similar distribution of units being more circular (*e*.*g*., a2 and b249) or more elongated (*e*.*g*., b241 and a35). However, this analysis considers only two of the four dimensions of connection weights and ignores the potential covariance structure of the afferent map. We now turn to the covariance structure of afferent connectivity from well-fit model units.

### 3.5 Dimensionality of the afferent field

To assess the structure of afferent tuning, we compute the expected covariance between each pair of the four tuning dimensions. This calculation is complicated by the differing coordinates across dimensions, with only horizontal and vertical space being comparable. Within limits imposed by its periodic nature, the orientation dimension can be represented in octaves, allowing a natural comparison to its spectral pair, *i*.*e*., spatial frequency (scale). To account for the extent of each receptive field in visual space, we normalize the spatial covariance of each model to have unit norm. This representation is unique up to an arbitrary scaling of the spectral and spatial dimensions. We choose this scaling empirically from our population data to equate the mean spectral and mean spatial covariance across all well-fit model units from V1 and V2.

Having established a common coordinate frame to examine the structure of the afferent weight envelope, we next consider the eigenstructure of the weight map’s four-dimensional covariance matrix. Qualitative inspection of the eigenvalue spectrum revealed units lying along a continuum between spherical (*i*.*e*., all eigenvalues of similar magnitude), and strongly elongated (*e*.*g*., dominated by a single principal eigenvalue). We quantify the *dimensionality* of an afferent map using the participation ratio of the eigenvalues *λ, i*.*e*.,

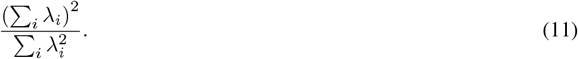

Fig. 7 depicts this dimensionality across populations, plotting the standard deviation of each afferent field in both scale and orientation (Fig. 7a,b). While we find the distribution of units across the V1 and V2 populations to be largely similar, we note in V2 multiple units of low dimensionality across a range of orientation bandwidths (*i*.*e*., cell a35 *vs*. cell a2). In contrast, V1 cells of large orientation bandwidth tend to have high dimensionality (*e*.*g*., a38, b241).

**Figure 7:**
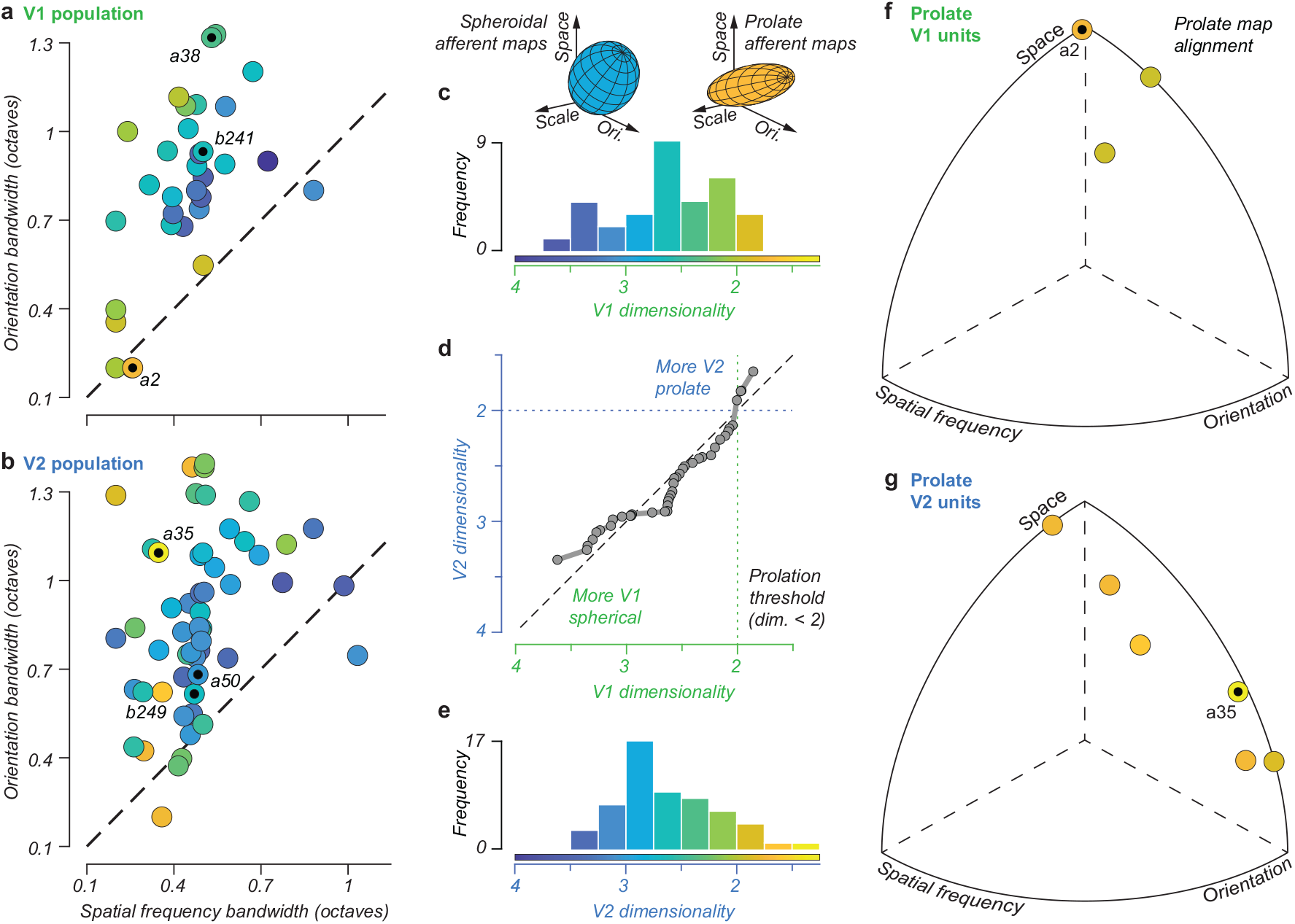
Covariance structure of model afferent weights. The standard deviation of afferent magnitudes in orientation and scale for units across our (a) V1 and (b) V2 populations, with color denoting (c,e) the dimensionality of each afferent field. (d) The cumulative distribution of afferent field dimensionality reveals that the most spherical units tend to be from V1, and most prolate tend to be from V2. (f,g) We visualize the alignment of prolate model units (defined as those having a dimensionality less than 2) as the direction of the principal axis over the covariance matrix of afferent weight magnitudes.

We may interpret example units’ dimensionality and spectral bandwidths by recalling the afferent field visualizations of Fig. 5b. V1 units are, in general, highly localized with (relatively) spheroidal afferent weight magnitudes (Fig. 7a, dimensionality > 2). An exception to this is the prolate cell a2, but a comparison with Fig. 7f reveals this unit to be spatially aligned, pooling vertical contrast over an elongated receptive field, co-linear with the preferred orientation. In contrast, V2 cells tend to be much more diffuse in the pooling of afferent activity, *e*.*g*., cell a35, which exhibits a linear receptive field tuned for spatially-displaced orthogonal orientations (see Fig. 5a). This receptive field arrangement is found to be prolate (Fig. 7f, dimensionality below 2) and aligned along the dimensions of both space and orientation, consistent with a curvature-tuned unit.

Interestingly, while the distribution of afferent dimensionality has a similar central tendency between V1 and V2 (Fig. 7c,e), a notable difference exists at the tails of each distribution: V1 has a higher frequency of units of high dimensionality, *i*.*e*., more spherical in afferent space, while V2 has more units that are more *prolate*, or elongated in a single dimension of afferent space (Fig. 7d). We also did not find units that were *oblate*, or flattened in two dimensions.

To investigate model unit prolation, we examine the alignment of the principal axis for units that are highly elliptical in afferent space (*i*.*e*., below a dimensionality of 2). We find prolate V1 units more aligned along spatial dimensions, having significant afferent weight covariance in space; *e*.*g*., cell a2, which has an elongated receptive field in the direction of its orientation tuning. In contrast, prolate V2 units have alignments spanning the orientation dimension, *e*.*g*., cell a35, which exhibits an afferent map highly correlated in space and orientation that is clearly evident in the linear component of its afferent field (Fig. 7g).

While in Fig. 3 we assess, to a limited extent, how well our model fits generalize to novel stimuli (*i*.*e*., a comparison of training and testing explained variance), we may instead measure how well models predict the neural response to stimulus classes beyond droplets. For this, we consider two additional data sets, gratings and textures, collected in addition to the droplet protocol for a subset of neurons (see methods). After fitting neurons to droplet data, models are fixed and exposed to grating and texture images matching experimental presentations. In Fig. 8a,c we examine observed and predicted tuning curves (averaged across phase) of example V1 and V2 cells. While some models appear to overestimate tuning bandwidth (*i*.*e*., cells a2 and a50), there is a strong qualitative agreement in preferred orientations and spatial frequencies. We similarly illustrate in Fig. 8b,d observed and predicted responses to textures, plotting the mean response (across instances and repetitions) of 5 synthetic texture families (naturalistic, dark) and their ‘scrambled’ spectrally-matched counterparts (noise, light).

**Figure 8:**
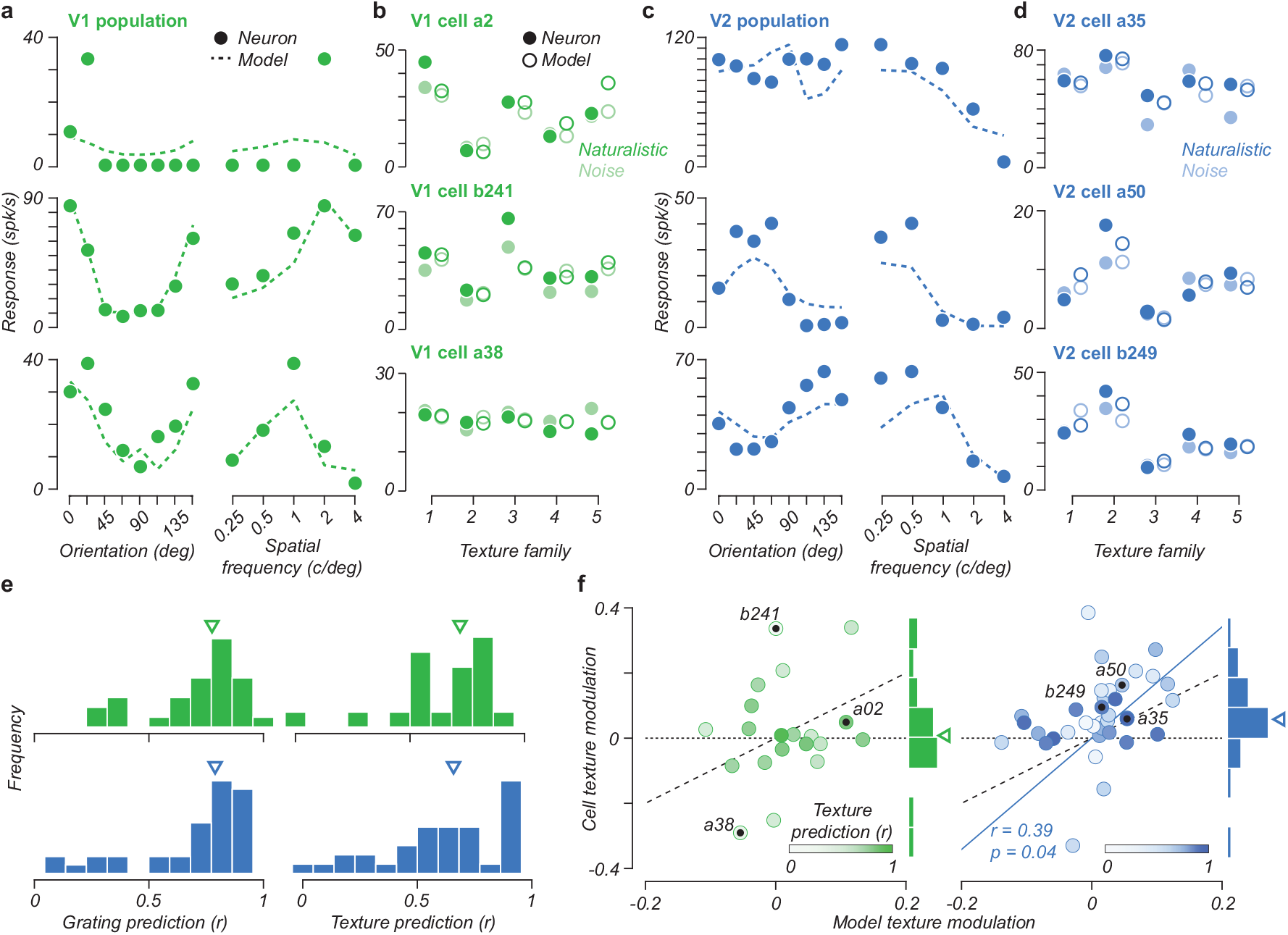
Comparisons of model predictions to recorded responses for oriented grating (a,c) and synthetic texture stimuli (b,d) for three examples of V1 (a,b) and V2 (c,d) neurons. Correlations (Pearson’s *r*) for grating stimuli are calculated after averaging responses over phase; for texture stimuli, responses are averaged over instances, correlating natural and noise stimuli across families. Grating/texture correlations of example cells were (top-down) .33/.83, .83/.28, .94/.52 for V1 and .80/.67, .85/.87, .84/.88 for V2. (e) Distributions of prediction accuracy. (f) Comparison of observed and predicted texture modulation across cells in our population, with saturation denoting texture prediction performance.

We assess prediction accuracy across the population by calculating the correlation (Pearson’s *r*) across responses, once for all gratings (Fig. 8a,c) and again for all textures (Fig. 8b,d), both naturalistic and noise; correlation histograms are given as Fig. 8e. Median prediction accuracy is high across V1 (*r* = 0.78, *r* = 0.72) and V2 (*r* = 0.79, *r* = 0.66), demonstrating our models generalize well across novel datasets. Importantly, we then measure the extent to which fitted model units capture *texture modulation, i*.*e*. a tuning for naturalistic textures over noise while controlling for local spectral content, as observed across recorded neurons. Fig. 8f depicts the texture modulation of each cell versus its fitted model. As expected, no significant texture modulation was observed in V1, while the population of V2 cells has a positive modulation index central tendency[12] (median of 0.07; *t*(35) = 3.37, *p* = 0.001), responding more strongly to naturalistic texture than spectrally-matched noise. Notably, many model units reproduce texture modulation exhibited by cells, *i*.*e*. cell a35, particularly true when excluding units of lower texture prediction quartile that poorly predict texture responses (*r* ≤ 0.45). Modulation indices of fitted models and cells are correlated (Fig. 8f blue line, *r* = 0.39, *p* = 0.040), demonstrating the afferent model’s ability to capture the origin of V2 selectivity for higher-order stimulus features not seen in V1.

It should be noted that the results presented are for a subset of model fits, *i*.*e*. those predicting more than 10% of variance from withheld (testing) data, that best reflect true neural tuning by well-generalizing across our recordings. However, While not shown here, this exclusion criterion (testing V.E. *<* 0.1) was explored (± 50%) and found to have little qualitative impact on our results. Outside of this range, the remaining population was too small to retain statistical power or had effects dominated by the overfitting of noisy cells. The collection of additional datasets will allow for this criterion to be relaxed: Fig. 3a,c demonstrate virtually all effectual recordings capture some predictive power.

## 4 Discussion

Primary visual cortex (area V1) has been intensively studied for more than sixty years. In primates, most V1 neurons are orientation selective, and their responses are well captured by models constructed by pooling the normalized and rectified responses of a small number of oriented filters [1, 17, 20, 22, 30, 33, 37, 42, 56]. Although receiving most of its afferent input from V1, an account of the responses of neurons in the second visual area (V2) has been more elusive. In many studies, V2 receptive field properties seem similar or even indistinguishable from those of V1[18]. The most robust reported differences are receptive field size (V2 RFs are roughly twice as large as those of V1), selectivity for figure-ground relationships (“border ownership”) [61], selectivity for relative, rather than absolute, binocular disparity [52], and selectivity for higher-order texture statistics [12, 60].

Here, we sought to characterize and distinguish cells in these two areas by fitting and comparing their responses within a common model. The first stage computes half-wave rectified responses of a fixed bank of oriented bandpass filters, mimicking a complete population of V1 simple cells. A second stage computes a weighted combination of these responses, followed by an output nonlinearity, to capture the firing rate of a single neuron. The second stage optimized against firing rates of individual V1 and V2 neurons, elicited by a novel stimulus ensemble consisting of random sparse and locally-oriented “droplets” designed to drive a (relatively) sparse subset of V1 afferents. We optimized the parameter likelihood given observed stimuli and firing rates, augmented by regularization terms to encourage sparsity and both spatial and spectral locality.

The fitted model provides a good account of the responses of cells well-driven by the droplet ensemble in both visual areas, matching or exceeding the performance of previous models[58]. Specifically, the model accounts for 38% of the explainable variance of V2 and 43% of V1 responses. The model also generalizes well to stimuli other than droplets, predicting responses to sinusoidal gratings and synthetic texture stimuli (Fig. 8). For comparison, a similarly-structured model, using a linear combination of rectified responses of a fixed bank of oriented binary-valued filters, accounted for 40% explainable variance of V1 and 30% of V2 over responses to high-contrast natural images [58]. Another LNL model, using quadratic nonlinearities, reports cross-correlation values for the model, fitted to natural image responses, in the range [0, 0.4][26].

Our fitted models provide some insights into the nature and origin of V2 response properties. Specifically, consistent with previous models [26, 58], we find that many V2 cells are V1-like, being well-described by the spatially pooled rectified responses of a *single* oriented convolutional filter. But some V2 cells are described by more complex combinations of afferents (Fig. 4b, 5, and 7) that involve inseparable combinations over orientation, scale, and position, consistent with previous findings[26]. Some neurons in V2 display novel selectivities not observed in V1, such as size-invariant chromatic selectivity [50], and relative disparity selectivity in three-dimensional scenes [52]. Anatomical studies illustrating the partition of V2 into different stripe compartments based on staining for the respiratory chain enzyme cytochrome oxidase [49], which suggests that there might be different functional clusters with distinct tuning [26, 27, 53]. It may be that the “V1-like” neurons we observe in V2 are simply selective for features more nonlinear than those supported by the model.

Unlike Willmore et al. 58, however, we did not find a significant increase of afferent suppression in V2 relative to V1. One possibility for this difference is that, due to our model’s inclusion of an output nonlinearity, fits could constrain response activity without additional suppressive connection weights. Further, we did not observe the elongated V2 receptive fields, *i*.*e*. those tuned to a single orientation and spatial frequency with a higher aspect ratio, reported by Liu et al. 26. These tuning characteristics may have been avoided during optimization by our spatial locality bias or missed in our analysis of prolate units covariance, which, by combining spatial dimensions, does not evaluate an afferent map’s aspect ratio.

While our model captures responses to commonly used stimulus families, its simplicity may restrict its ability to reveal more complex nonlinear properties. In particular, despite the successful prediction of responses across different types of texture distinguished by their orientation and spatial frequency content (Fig. 8e), the model largely fails to capture the substantial modulation of V2 neurons by naturalistic vs. noise texture stimulation (Fig. 8f, [12]). The higher-order statistical structure of texture models [36] as well as preliminary model exploration, suggests that this property can emerge naturally from the inclusion of pooling in the second stage of the model (*i*.*e*., V2 “complex cells”) [62]. The first stage of the model is also oversimplified, deliberately omitting known properties of V1 simple cells. Specifically, the receptive fields are idealized, with a single bandwidth (one octave) and orientation tuning (60 deg) and a discrete set of preferred orientations and scales, while V1 simple cells exhibit diversity in these respects. In our model, the first-stage nonlinearity is half-wave rectification, whereas V1 simple cells are better approximated by half-squaring [16, 42], and again exhibit diversity. Finally, V1 neurons are suppressed by various stimuli, a property well captured by divisive normalization by a pool of neighboring cells [6, 17]. We leave the fitting of a more elaborate population of V1-like first-layer units to explain many, perhaps simultaneously recorded, V2 neurons as an intriguing avenue of future research. Nevertheless, the limitations and restrictions imposed on the current model were deliberate: Including additional stages or parameters would further complicate the optimization and interpretation of an already complicated model.

We offer the current model as an important, if incomplete, step toward building a model of V2 that has the richness and predictive power that V1 models now enjoy. Extending the model may require advances in model fitting, neural recording, and stimulus generation. However, even in the absence of these extensions, we are confident that this work moves us toward a clear and concise description of neural selectivity within the early visual cortex.

## 5 Acknowledgements

We are grateful to Kaitlyn Holman, Rui Pacheco, and Sullivan Bacerdo for their assistance, and to Manu Raghavan and Najib Majaj for their help with surgery, hardware, and software. This work was supported by grants from the National Institutes of Health (EY022428) and the Simons Foundation (543019) to J.A.M. and E.P.S. J.D.L was supported in part by a Leon Levy Fellowship in Neuroscience.

